# Removal of a subset of non-essential genes fully attenuates a highly virulent *Mycoplasma* strain

**DOI:** 10.1101/508978

**Authors:** Joerg Jores, Li Ma, Paul Ssajjakambwe, Elise Schieck, Anne Liljander, Suchismita Chandran, Michael Stoffel, Valentina Cippa, Yonathan Arfi, Nacyra Asad-Garcia, Laurent Falquet, Pascal Sirand-Pugnet, Alain Blanchard, Carole Lartigue, Horst Posthaus, Fabien Labroussaa, Sanjay Vashee

## Abstract

Mycoplasmas are the smallest free-living organisms and cause a number of economically important diseases affecting humans, animals, insects and plants. Here, we demonstrate that highly virulent *Mycoplasma mycoides* subspecies *capri* (Mmc) can be fully attenuated *via* targeted deletion of non-essential genes encoding, among others, potential virulence traits. Five genomic regions, representing approximately ten percent of the original *Mmc* genome, were successively deleted using *Saccharomyces cerevisiae* as an engineering platform. Specifically, a total of 68 genes out of the 432 genes verified to be individually nonessential in the JCVI-Syn3.0 minimal cell, were excised from the genome. *In vitro* characterization showed that this mutant was similar to its parental strain in terms of its doubling time, even though ten percent of the genome content were removed. A novel *in vivo* challenge model in goats revealed that the wild-type parental strain caused marked necrotizing inflammation at the site of inoculation, septicemia and all animals reaching endpoint criteria within seven days after experimental infection. This is in contrast to the mutant strain, which caused no clinical signs nor pathomorphological lesions. These results highlight, for the first time, the rational design, construction and complete attenuation of a *Mycoplasma* strain via synthetic genomics tools. Trait addition using the yeast-based genome engineering platform and subsequent *in vitro* or *in vivo* trials employing the *Mycoplasma* chassis will allow us to dissect the role of individual candidate *Mycoplasma* virulence factors and lead the way for the development of an attenuated designer vaccine.

**IMPORTANCE:** Members of the *Mycoplasma mycoides* cluster cause important animal plaques in Africa and Asia, which impact animal welfare, provision of food and the life of thousands of small-scale farmers. We applied synthetic biology tools to *Mycoplasma mycoides* in order to design and create a fully attenuated *Mycoplasma* strain that was subsequently confirmed *in vivo* using a novel caprine infection model. This is the first time that a *Mycoplasma* mutant developed by applying synthetic biology tools has been tested *in vivo* to pin point candidate virulence traits. The mutant strain is similar to “apathogenic *E. coli* K12” strains that boosted the research on host-pathogen interactions for the genus *Escherichia* and other bacterial genera.

## INTRODUCTION

Bacteria belonging to the genus *Mycoplasma* are wall-less bacteria that cause massive economic losses in the livestock sector (chickens, ruminants and pigs) and are responsible for human pneumonia and sexually transmitted diseases (STDs). Currently, there is an absence of commercial vaccines against infections with the human pathogens *Mycoplasma pneumoniae* and *Mycoplasma genitalium* (1). In contrast, many livestock vaccines are commercialized, which rely either on adjuvanted killed bacteria or on attenuated strains obtained after successive rounds of sub-culturing or chemical mutagenesis (2). Due to these empirical approaches, the exact mechanism triggering the attenuation is unknown for many of the previously developed live attenuated *Mycoplasma* vaccines. Strikingly, these vaccines are far from being optimal since they often display short durations of immunity and limited efficacy (3-5). A better understanding of pathogenicity and the identification of virulence traits would foster next generation vaccines.

For many years, the lack of genetic tools has limited our basic understanding of *Mycoplasma* pathogenicity. Due to their regressive evolution by gene loss, mycoplasmas appear to lack many of the common bacterial effectors and toxins used to interact with their hosts or to escape the hosts’ immune systems (6, 7). Lipoproteins have been proposed to be involved in both aspects by using their cytoadherent properties and allowing antigenic variability through phase or sequence variation (8). Other candidate virulence traits, such as the *Mycoplasma* Ig binding *protein-Mycoplasma* Ig protease (MIB-MIP) system (9) and the hydrogen peroxide production system (10) have been suggested, but not confirmed *in vivo.*

The availability of a genome engineering platform that allows directed and precise mutagenesis for *Mycoplasma mycoides* is undoubtedly a new starting point towards better understanding of host-pathogen interactions. In this work, we engineered a *Mmc* strain by deleting approximately one tenth of the genome, including candidate virulence traits. The resulting mutant retains almost wild-type like growth characteristics and was attenuated both *in vitro* and *in vivo.* The construction of this fully attenuated and safe laboratory *Mycoplasma* strain paves the way for research into host-pathogen interactions and is a good starting point to revisit the actual role of suggested virulence determinants in *Mycoplasma*.

## RESULTS

### Generation of the mutant strain GM12::YCpMmyc1.1-*∆68*

To demonstrate attenuation of *Mmc* by rational design, five genomic regions were targeted in this study. The precise localizations in the *Mmc* GM12 genome are shown in Figure 1. The first two target deletion regions contained genes encoding the glycerol-dependent hydrogen peroxide metabolic pathway and its suggested ABC transporter encoded by the *gtsABCD* operon (11). This pathway has been suggested to be a main virulence mechanism for *M. mycoides* (11), but *in vivo* confirmation is still missing and in *M. gallisepticum* the pathway does not seem to be linked to virulence (12). Thus, the genes *glpF, glpK* and *glpO* (MMCAP2_0217-0219; 2,984-bp region; D1) and the *gts* gene region that includes the gene *lppB* (MMCAP2_0456-0459; 4,950-bp region; D2) were deleted in the *Mmc* genome by the yeast-based engineering method (13). As previously mentioned, lipoproteins were another target of interest since they likely trigger not only host-pathogen interactions but also, overwhelming immune reactions that result in inflammation (14). The lipoproteins encoded in the D3 (MMCAP2_0014-0016; 4,677-bp region; D3) as well as the D5 region (*lppQ,* MMCAP2_0889-0904; 24,906-bp region, D5) were also excised employing again the yeast-based engineering method. Finally, we deleted a large genomic region that contained the *Mycoplasma-specific* F1-likeX0 ATPase (15), the MIB-MIP system (9) that has been shown *in vitro* to modulate the action of immunoglobulins through specific degradation, an integrative and conjugative element (ICE) (16) and several genes encoding other lipoproteins. The ICE was targeted in an effort to reduce mobile elements from the *Mmc* genome. In this case, about 70 kbp (MMCAP2_0550-0591; 69,220-bp region; D4) were targeted and deleted from the *Mmc* genome using the yeast-based engineering method in one stretch.

**Figure 1:**
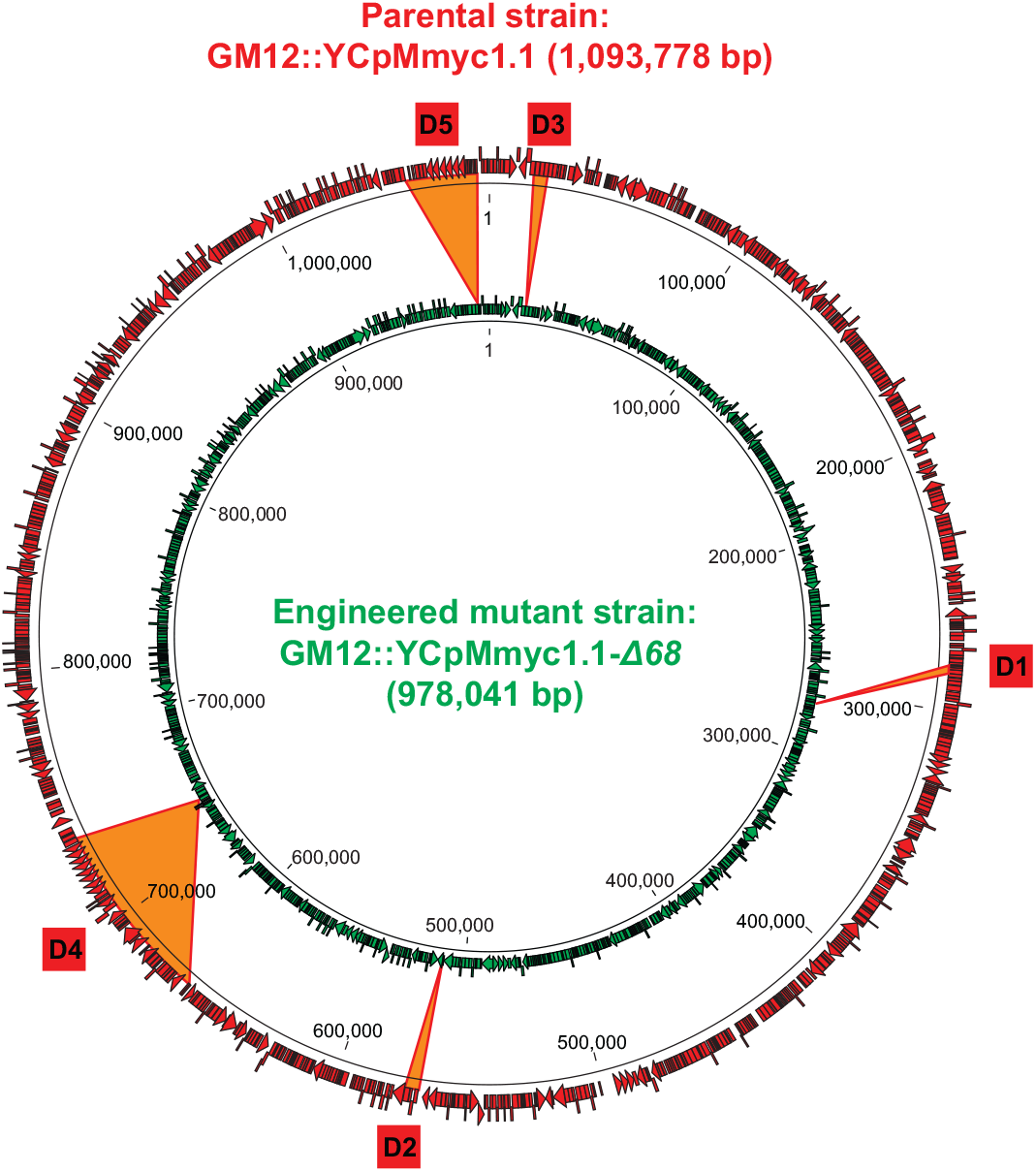
Design of deletion mutant *Mmc* strain. (A) Schematic displaying the genomes of the isogenic parental strain GM12::YCpMmyc1.1 and its derivative, the deletion mutant GM12::YCpMmyc1.1-*∆68*.

After each cycle of deletions, the modified *Mmc* genome was isolated from yeast cells and transplanted back into *M. capricolum* subsp. *capricolum* (*Mcc*) recipient cells to confirm the viability of each mutant *Mmc* strain. Overall, the final mutant strain, named *Mmc* GM12::YCpMmyc1.1-*∆68*, was generated in five sequential deletion cycles (Figure 1). The gene knock-outs were verified by amplifying across each deleted region (Figure S1). Genomic DNA from the GM12::YCpMmyc1.1-*∆68* was isolated and analyzed by sequencing to confirm the deletions (Figure1B). The genome sequence of GM12::YCpMmyc1.1-*∆68* was deposited at the ENA database under the accession number LS483503.

### The mutant strain GM12::YCpMmyc1.1-*∆68* is viable and unaffected in its morphology or growth in axenic medium

The colonies of *Mmc* GM12::YCpMmyc1.1-*∆68* were of similar size to those of GM12::YCpMmyc1.1 and GM12. Cell morphology of the GM12, the isogenic parental strain GM12::YCpMmyc1.1 and GM12::YCpMmyc1.1-*∆68* strains was evaluated using scanning electron microscopy (Figure 2A). All strains tested were globular in shape and lacked any special morphological features. The diameter of the microorganisms was in the range of 500 nm, as expected for a *Mycoplasma* cell. The mutant GM12::YCpMmyc1.1-*∆68* grew with a doubling time somewhat similar to that of the parental strains GM12 and GM12::YCpMmyc1.1 (Figure 2B). Together, these results strongly suggest that the deletion of approximately 100 kbp of genomic content from the *Mmc* genome did not adversely affect structural integrity or *in vitro* growth of the mutant GM12::YCpMmyc1.1-*∆68*.

**Figure 2:**
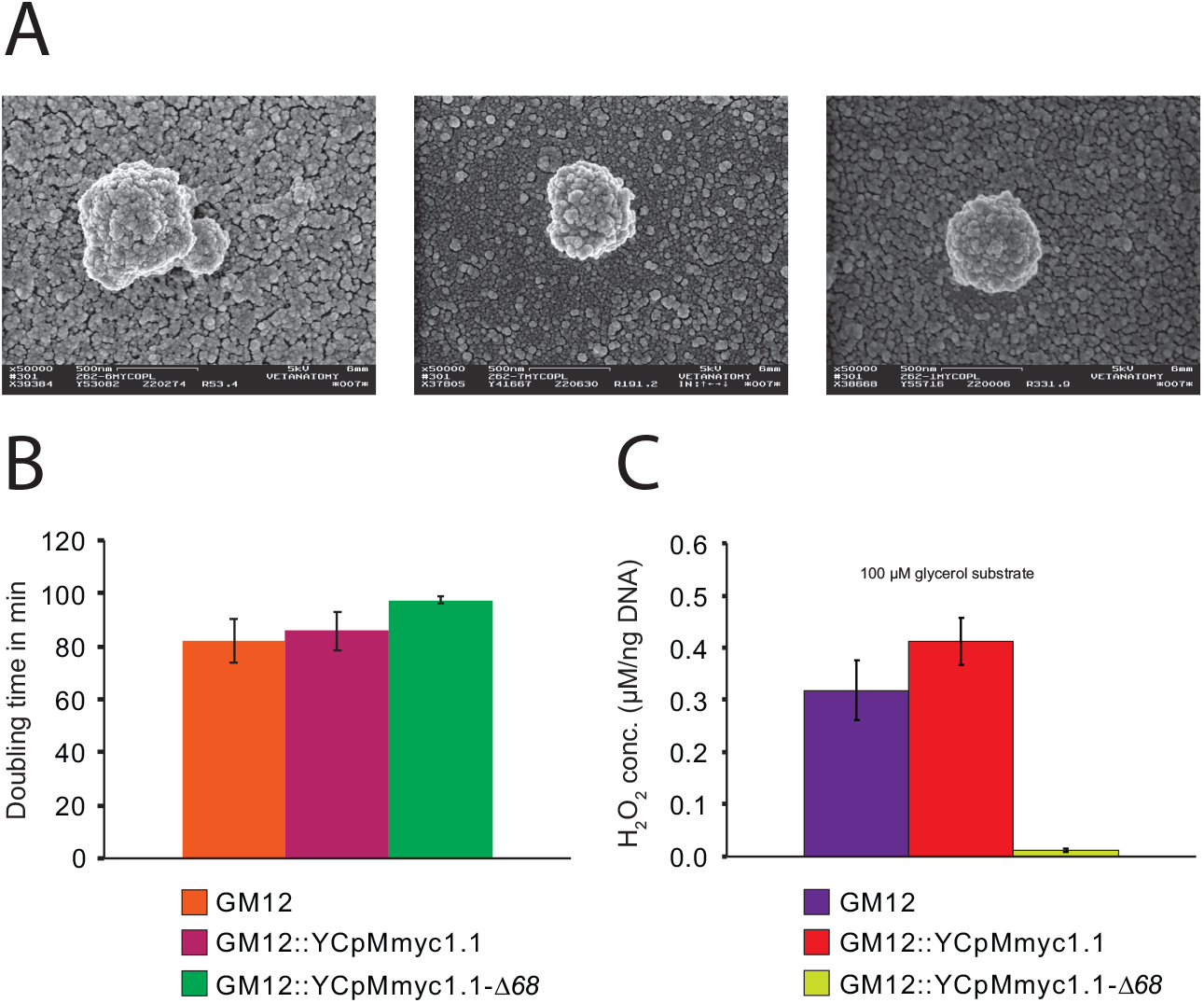
*In vitro* characteristics of the parental strains GM12, GM12::YCpMmyc1.1 and its deletion mutant GM12::YCpMmyc1.1-Δ*68* with respect to their morphology revealed by scanning electron microscopy (A), their doubling time (B), their production of hydrogen peroxide in the presence of glycerol (C), the bars display the standard deviation in B and C

### Inability of the mutant strain GM12::YCpMmyc1.1-*∆68* to produce hydrogen peroxide in the presence of glycerol *in vitro*

This pathway was completely deleted in the construction of the mutant strain GM12::YCpMmyc1.1-*∆68*. Therefore to phenotypically confirm the deletion, we measured and compared hydrogen peroxide production levels between the control GM12, GM12::YCpMmyc1.1 and GM12::YCpMmyc1.1-*∆68 in vitro.* In the presence of the glycerol substrate, the mutant GM12::YCpMmyc1.1-*∆68* shows a significant decrease in hydrogen peroxide production when compared to its parental strains (Figure 2C). Indeed, while GM12 and GM12::YCpMmyc1.1 produced >0.3 μM of H_2_O_2_, the mutant strain produced very low amounts of H_2_O_2_ (0.01 μM), at least 30-fold lower under these conditions.

### Inability of the mutant strain GM12::YCpMmyc1.1-*∆68* to degrade immunoglobulin *in vitro*

Another potential virulence trait encoded by mycoplasmas is the MIB-MIP system, which may play a role in immune evasion by cleavage of immunoglobulins (Figure 3A) (9). Incubation of caprine IgG with GM12, GM 12::YCpMmyc 1.1 and GM 12::YCpMmyc1.1-*Δ68* showed a clear difference in the strains’ abilities to degrade IgG (Figure 3B). The two bands at 25 kDa and 50kDa corresponds to the IgG light and heavy chains. The mutant strain GM 12::Y CpMmyc 1.1-*∆68* exhibited no degradation of IgG, as noted by the lack of the 44 kDa band (Figure 3B, black asterisk). This band, clearly visible in the other two strains, is indicative of proteolytic cleavage of the IgG heavy chain. Another pattern of degradation, with a band at a size of about 30 kDa, is visible in the three strains. It was previously reported that this IgG cleavage is not specific or directly linked to the MIB-MIP system (9).

**Figure 3:**
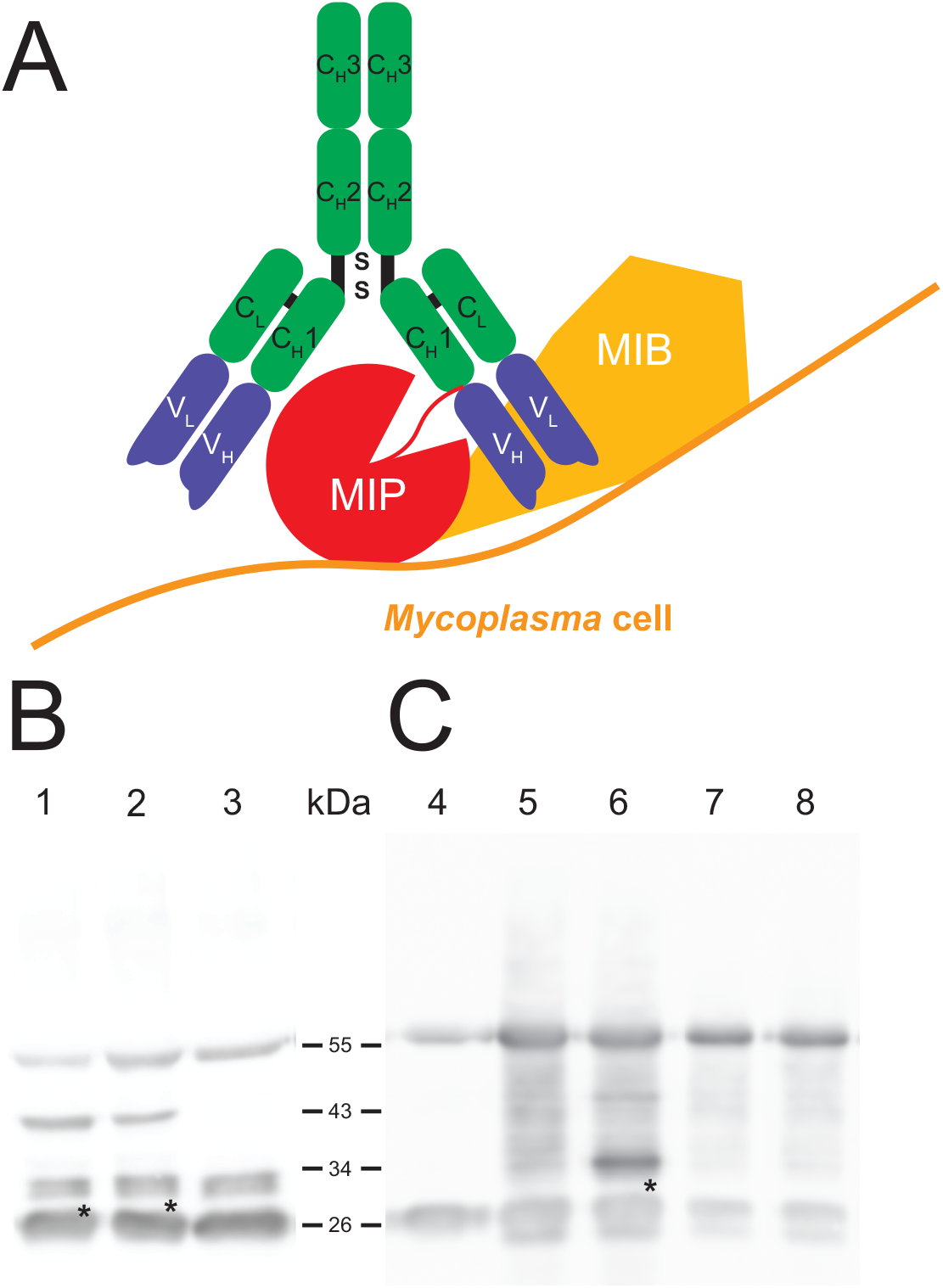
*In vitro* and *in vivo* ability of the *Mmc* MIB-MIP system to degrade caprine immunoglobulin: (A) cartoon displaying the suggested action of the MIB-MIP system, (B) *in vitro* ability of the parental strains GM12, GM12::YCpMmyc1.1 and their deletion mutant GM12::YCpMmyc1.1-Δ*68* to degrade caprine immunoglobulin, 1-GM12 + 5 μg caprine IgG, 2-GM12::YCpMmyc1.1 + 5 μg caprine IgG, 3-GM12::YCpMmyc1.1-Δ*68* + 5 μg caprine IgG; the black asterisk marks the 44 kDa cleaved fragment (C) *In vivo* detection of degradation of the *Mmc* MIB-MIP system using serum samples from the animal CK51 that succumbed from disease and had septicemia, 4-5 μg caprine IgG, 5-serum from goat CK51: −1 dpi (GM12 group), 6-serum from goat CK51: 4 dpi (GM12 group), 7-serum from goat CK45:-1 dpi (GM12::YCpMmyc1.1-Δ*68* group), 8-serum from goat CK45: 4 dpi (GM12::YCpMmyc1.1-Δ*68* group)

### The mutant strain GM12::YCpMmyc1.1-*∆68* is fully attenuated *in vivo*

We next tested whether GM12::YCpMmyc1.1-*∆68* was able to cause disease in its native host. Sixteen male outbred goats (*Capra aegagrus hircus*) were used in this animal infection trial. The animals were separated into two groups of equal numbers. After the infection, no immediate clinical signs of disease were observed. Two animals in the GM12::YCpMmyc1.1-*∆68* group had to be removed from the experiment, because of acquired wounds unrelated to the infectious agent. Animal euthanasia was planned 28 days post infection (dpi). However, all eight animals inoculated with the GM12 strain developed severe clinical signs, with pyrexia starting 2-3 dpi (Figure 4B). Their body temperature continued to increase, up to 41-41.5°C, during the following days. The animals stopped feeding, were apathic and showed signs of pain. According to the endpoint criteria stated in the animal experiment protocol, they had to be euthanized between 5-6 dpi (Figure 4A, red line). In sharp contrast, animals inoculated with the GM12::YCpMmyc1.1-*∆68* mutant strain did not develop any clinical signs of disease and were all monitored until the end of the trial (Figure 4A). Their body temperatures fluctuated between 38°C and 39°C, with a few isolated cases where animals showed temperatures above 39°C but, never for more than 2 consecutive days (Figure 4B). The animals appeared to remain healthy and gained weight during the experiment (Figure 4C). Their heartbeat and respiratory rates, between 80-110 beats/min and 20-30 breaths/min, respectively, remained constant over the course of experimentation.

**Figure 4:**
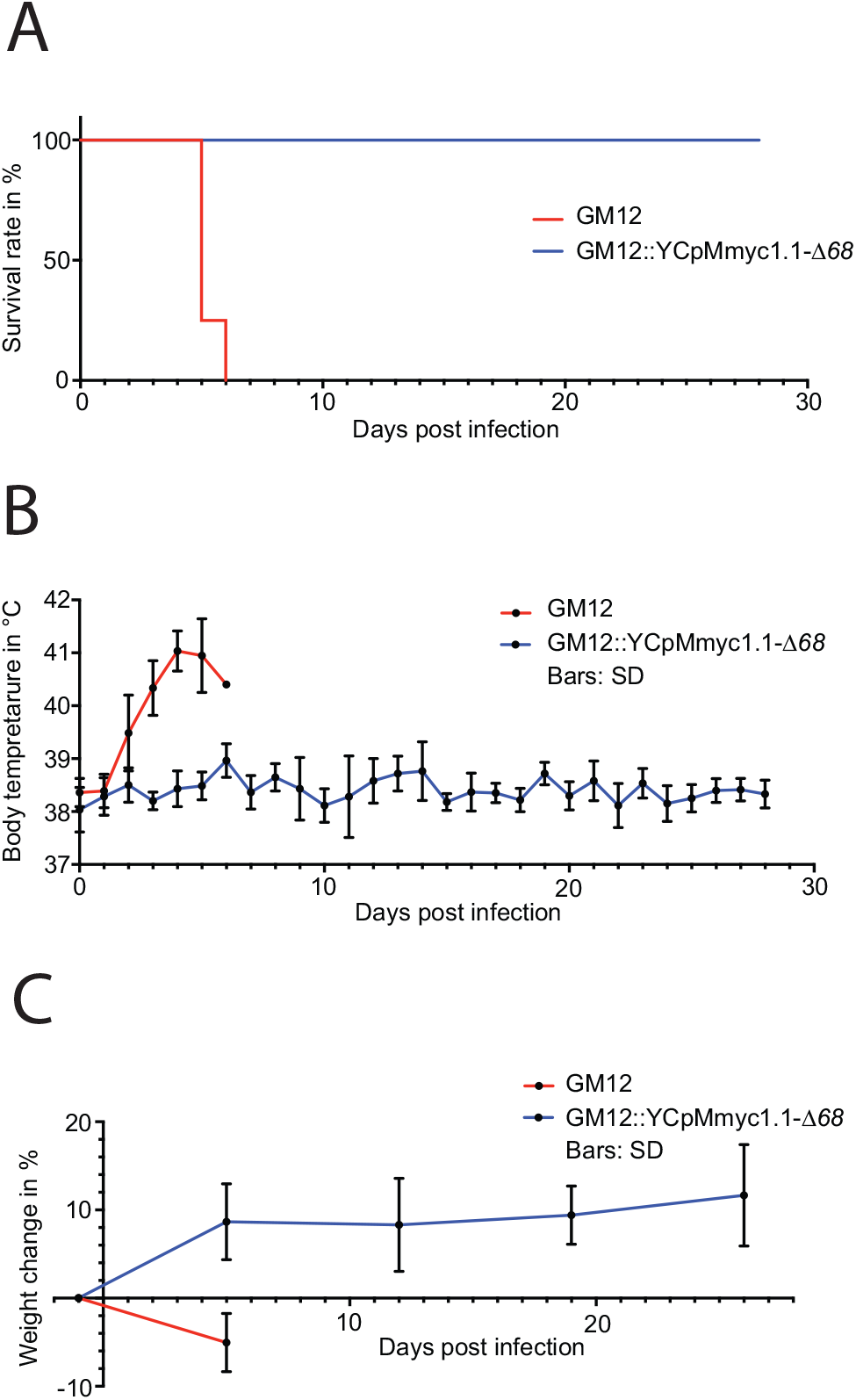
Comparison of the clinical parameters monitored during the in vivo challenge trial between the animals that received the GM12 and its derivative GM12::YCpMmyc1.1-Δ*68*. (A) Kaplan Meier survival curve based on animals reaching endpoint criteria, (B) Average body temperatures during experimental infection. Values were generated using daily rectal temperatures from the two groups. (C) Average weight gain during experimental infection. Values were generated using interval measures from the two groups. The standard deviations are displayed as bars in B and C.

In all animals infected with the GM12 strain, the main pathological lesions were similar, with a severe and extensive inflammation of the soft tissues of the neck around the site of inoculation. Additional macroscopic findings were severe pulmonary edema and congestion. Histologically, there was extensive coagulation necrosis of the connective tissue (Figure 5B, thick arrows) and musculature surrounding the trachea, in the vicinity of the inoculation site (Figure 5B and D, diamonds). A marked infiltration of mainly degenerate neutrophilic granulocytes (Figure 5B and D, asterisks) was always found associated with the necrosis. The necrotizing process extended to the trachea, the subcutis and skin. In addition, all animals showed multifocal acute necrosis with infiltration of neutrophilic granulocytes in liver and kidney and for three animals, in the lung. These lesions were indicative of an acute septicemia. Among all animals infected with the mutant strain GM12::YCpMmyc1.1-Δ*68*, and upon euthanasia 28-29 dpi, no pathological lesions were found around the inoculation site. The soft tissue around the inoculation site was within normal limits. Additionally, neither inflammation nor necrosis associated with the epithelium, the submucosa or any cartilage tissues was histologically observed (Figure 5, panels A and C).

**Figure 5:**
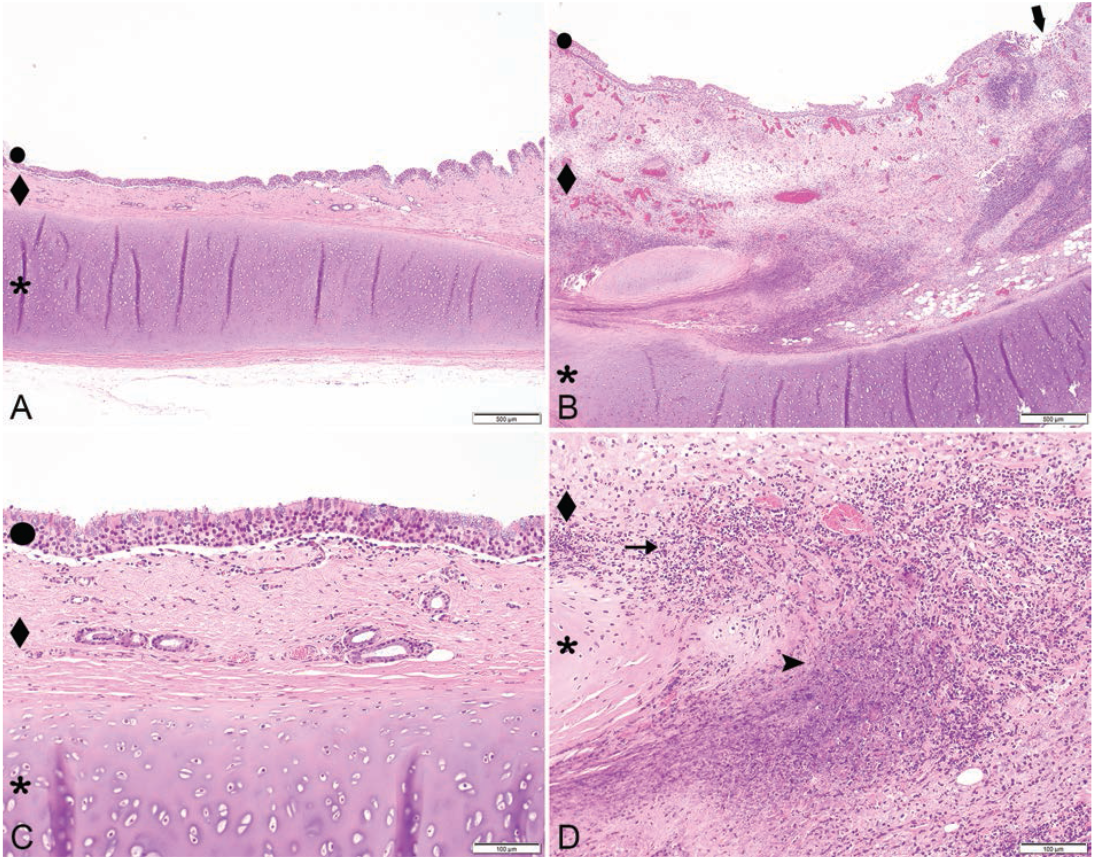
Composite figure displaying representative histological results from tracheal tissue at the inoculation site. Tissues were stained with hematoxylin and eosin. Low (40x) and high (200x) magnification of trachea of a goat inoculated with *Mmc* GM12::YCpMmyc1.1-Δ*68* (panels A and C) depicting unaffected epithelium (circle), submucosa (diamond) and cartilage (asterics). or of a goat inoculated with *Mmc* GM12 (panels B and D), depicting ulceration of the epithelium (thick arrow, panel B), massive extension of the submucosa due to extensive areas of necrosis (diamond, panels B and D) and infiltration with large numbers of degenerate neutrophilic granulocytes (asterisk, panels B and D). Size standards are displayed in each picture.

*Mmc* GM12 was re-isolated from the blood of all animals experimentally infected with the wild-type parental strain. Bacteremia was characterized by 10^6^ up to 10^9^ CCU.ml^-1^ of blood, as measured by serial dilutions (Table 1). Blood samples collected from goats infected with GM12::YCpMmyc1.1-Δ*68* did not reveal any bacteremia.

**Table 1:** Summary of post mortem records of goats.

| Animal No. | Date of euthanasia | Bacteremia ccu/ml | Inflammation of the neck around the injection site | Pulmonary congestion | Pulmonary oedema | Congested kidneys | Mucoid enteritis & congestion | Pleural fluid in thoracic cavity | Adherent lung to rib cage | Liver abscess |
| --- | --- | --- | --- | --- | --- | --- | --- | --- | --- | --- |
| CK032 | 6 dpi | 10<sup>7</sup> | X | X | X | X | X | X |  |  |
| CK034 | 5 dpi | 10<sup>7</sup> | X | X | X | X | X |  |  |  |
| **CK035** | 29 dpi |  |  |  |  |  |  |  |  |  |
| CK040 | 5 dpi | 10<sup>8</sup> | X | X | X | X | X |  |  | X |
| CK043 | 5 dpi | 10<sup>8</sup> | X | X | X | X | X |  |  |  |
| **CK045** | 28 dpi |  |  |  |  |  |  |  |  |  |
| CK046 | 5 dpi | 10<sup>8</sup> | X | X | X | X | X |  |  |  |
| **CK047** | 29 dpi |  |  |  |  |  |  |  |  |  |
| CK048 | 5 dpi | 10<sup>9</sup> | X | X | X | X | X |  |  |  |
| **CK049** | 28 dpi |  |  |  |  |  |  |  |  |  |
| CK051 | 5 dpi | 10<sup>9</sup> | X | X | X | X | X |  |  |  |
| **CL001** | 28 dpi |  |  |  |  |  |  |  |  |  |
| CL002 | 5 dpi | 10<sup>8</sup> | X | X | X | X | X |  | X |  |
| **CL003** | 28 dpi |  |  |  |  |  |  |  |  |  |

Animals displayed in bold were infected with GM12::YCpMmyc1.1-Δ68, the other animals were infected with wtGM12; dpi - days post infection,
ccu – color changing unit

### The MIB-MIP system in *Mmc* GM12 is functional *in vivo*

The MIB-MIP system was shown to be active *in vitro,* and to be present in large amounts at the cell surface (17) during infection (18). Two animal sera were selected from the *in vivo* infection ‡ trial: CK51 from the GM12 group and CK45 from the GM12::YCpMmyc1.1-*∆68* group and tested for cleavage of IgG. The pre-infection sera from both animals demonstrated no proteolytic cleavage of IgG when compared to the IgG control (Figure 3C). Conversely, the post-infection serum of CK51, which had succumbed to disease and had a high titer of bacteria Š in the venous blood, clearly exhibited the typical band at 44 kDa, consistent with the size of a cleaved IgG heavy chain by the MIB-MIP system. As expected, no band at 44 kDa was seen in the post-infection serum of CK45. This clearly demonstrates, for the first time, that the MIB-MIP system is functional within the caprine host and that its proteolytic activity is triggered during an *Mmc* infection.

## DISCUSSION

The first aim of this work was to fully attenuate a highly pathogenic strain of *Mycoplasma mycoides* following a rational deletion design. The second aim was to verify this attenuation *in vivo* using the native host, since no rodent animal models for highly virulent *M. mycoides* exist (3). Many candidate virulence factors of *Mycoplasma mycoides* have been suggested but, all except capsular polysaccharide (19) have never been confirmed according to Falkow’s postulates *in vivo* (20).

First, we relied on previous knowledge to select five genomic regions that encode candidate virulence traits. These regions, distributed around the *Mmc* genome, consisted of 68 genes. The first two regions (D1 and D2) encode enzymes and putative glycerol transporters involved in the production of hydrogen peroxide using the glycerol-dependent metabolism of mycoplasmas (10). The region D3 encodes major antigens (LppA/P72) in *Mycoplasma mycoides* (21) that induced T cell responses early in infection (22). The region D4 includes an integrative conjugative element (ICE), the MIB-MIP system (9) and the F1-likeX0 ATPase (15). The last region (D5) encodes six lipoproteins including the lipoprotein Q (LppQ), which has been suspected to be involved in exacerbating immune responses and other virulence determinants (23). Interestingly, among these 68 genes deleted in our mutant, 67 were defined as non-essential in the JCVI-Syn3.0 minimal cell, a study in which 432 genes were classified as non-essential to sustain the life of a minimal *Mycoplasma* cell (24). Only one gene encoding the glycerol phosphate kinase (GlpK) (MMCAP2_0218, region D1), which was retained in the minimal JCVI-Syn3.0 cell but, classified as a “quasi-essential” gene. This means that the function encoded by the glycerol kinase, i.e. the transfer of a phosphate group on the glycerol molecule to produce glycerol-3-phosphate, could be compensated for by another gene which encodes a similar function in the full-length genome. The compensatory gene for *glpK* is probably absent in the minimal JCVI-Syn3.0 cell, but present in our mutant GM12::YCpMmyc1.1-Δ*68*, allowing the deletion of *glpK* without any defect in growth. Therefore, our results are consistent with the quasi-essentiality of *glpK.* Altogether we retained 69 out of the 87 lipoproteins (D1-0, D2-1, D3-3, D4-8, D5-6) present in GM12, while the minimal cell retained only 15 lipoproteins (24).

It was paramount for us that the mutant strain GM12::YCpMmyc1.1-Δ*68* maintains a doubling time similar to its parental strain, since we wanted to create a ‘K12-like’ strain that can be further used as a cellular platform to introduce antigens or stretches of DNA. It was interesting for us to observe that, despite complete removal of the glycerol pathway, which maybe important for cell metabolism, there was no significant impact on *in vitro* growth. It is known that there is a trade-off between genome size and growth rate. The drastic deletions in the genome of JCVI-syn3.0 strain led to a substantial increase in the generation time, from ~60 min to ~180 min (24). Recently, we also have shown that the deletion of a gene encoding an enzyme important for synthesis of carbohydrates can subsequently lead to an increase in the generation time (25). However, in this work, we significantly reduced the genome of GM12 by more than 100 kbp (i.e. 106,737 bp) without seeing any significant difference in the growth rate of GM12::YCpMmyc1.1-Δ*68* in comparison to the wild-type strain. Still, this reduction represents ~10% of the initial genome size confirming that, in addition to the size of the deletions, the nature of the genes deleted is also very likely to influence the generation time of mycoplasmas. The main goal of this work was to construct a fully attenuated *Mmc* strain, that is safe to handle in the laboratory. The first confirmation of the attenuation of the GM12::YCpMmyc1.1-Δ*68* strain was obtained *in vitro.* The production of hydrogen peroxide was almost completely abolished in the mutant strain, confirming the participation of the *glpFKO* and/or *gtsABCD* pathways in the metabolism of glycerol. In addition, the loss of the IgG specific cleavage band at 44 kDa in GM12::YCpMmyc1.1-Δ*68* confirmed the role of the MIB-MIP system in the degradation of the host immunoglobulins.

To confirm the mutant strain’s attenuation *in vivo,* we developed an animal challenge model using Kenyan goats, outbred animals derived from different herds. The use of such animals increased variability to get a better idea of reproducibility and significance of the results (26). Animals infected with the GM12 strain developed specific symptoms (fever, heavy breathing, septicemia, etc.) and were all euthanized by 6 dpi. Strikingly, none of the goats infected with GM12::YCpMmyc1.1-*Δ68* developed such symptoms and were healthy for the entire course of the experimentation. These results exceeded our expectations and confirmed the complete abolishment of pathogenicty of the mutant strain GM12::YCpMmyc1.1-*Δ68*. In addition, the massive septicemia associated with very high titers of *Mycoplasma* observed in animals infected with the GM12 strain prompted us to investigate whether the MIB-MIP system would leave signatures of its action on immunoglobulins (Ig) in the serum of an animal (CK51) that had a titer of 10^9^ CFU/ml. Specific IgG cleavage, characteristic of the MIB-MIP system (9), was observed. No such cleavage was observed in the serum of animals infected with the mutant strain. This work shows, for the first time, that the MIB-MIP system of *Mycoplasma mycoides* is functional *in vivo. Mycoplasmas* have been viewed as stealth pathogenic organisms because they lack most of the immune activators or PAMPs found in other bacteria (27). Indeed, the lack of a cell wall or the capacity to produce either LPS or flagellins likely contribute to the chronicity of infection. The only PAMP that has been described for several *Mycoplasma* species is the surface lipoproteins, abundant components of their membrane (8). In the present study, we suggest another mechanism that could contribute to activating the immune system. Indeed, Ig cleaved by several bacteria, including those generated by *Mycoplasma hyorhinis,* have been described as ligands of the innate immune receptor LILRA2 (28). Once bound to this receptor, it triggers the activation of the innate immune system. It is also possible that this cleavage is in line with the ‘nutritional virulence’ of the parasite (29). The exact significance of the Ig cleavage regarding *Mycoplasma* infection of mucosal surfaces remains to be studied.

Interestingly, we observed severe inflammation around the site of injection in the animals that received GM12 whereas animals that were injected with the strain GM12::YCpMmyc1.1-Δ*68* developed no such pathomorphological lesions. Overwhelming immune reactions at the site of vaccination have been reported from immunizations against contagious bovine pleuropneumonia using live *M. mycoides* subsp. *mycoides* based vaccines such as T1/44, which is the closest relative of *Mmc* (30). Therefore, it is likely that any one or several of the deleted genes encode proteins that drive this overwhelming immune reaction in the GM12.

To conclude, we confirmed, *in vitro* and *in vivo*, our ability to design a fully attenuated strain *via* the precise reduction of ~10 % of the *Mmc* genome. However, we cannot currently pinpoint the weight of each deletion on the observed attenuation. The total clearance of the pathogen and the absence of a compelling humoral immune response, even at the inoculation point, is surprising and supports the total abolishment of pathogenicity. Now it is necessary to test more defined mutants such as a *glpOKF* mutant strain to get clarity about its real role in pathogenicity.

In addition, the design of next generation vaccines for, but not restricted to, *Mycoplasma* diseases will benefit from this study since a chassis that is fully attenuated and able to accommodate antigens for vaccine delivery that can be constructed based on our deletion mutant is now within reach. To induce a proper immune response via such a chassis, we have the option to add genes that appropriately stimulate an inflammatory immune response or alternatively, we can construct different chassis that direct responses towards Th1 or Th2 using TLR agonists. We consider a genetically modified *Mycoplasma* less problematic than other potential chassis since the survival time of *Mycoplasma* in general in the environment is very short. In addition, the unconventional codon usage (where UGA encodes tryptophan) and high AT content of *Mycoplasma* minimizes the risk of spread of genes to other bacteria. Additional experiments are necessary to decipher the role of individual virulence traits to understand these minimal bacterial pathogens better and to develop next generation rationale vaccines. Regardless, this study provides an attractive blueprint towards these goals, especially for those that are needed in low and middle-income countries.

## MATERIALS AND METHODS

### *Mycoplasma* strains

The *Mycoplasma mycoides* subsp. *capri* outbreak strain GM12 was used as positive control in the *in vivo* experiment (31). A modified *Mycoplasma capricolum* subsp. *capricolum* strain CK was used as recipient strain in genome transplantation protocols (13).

### Yeast strain and media

The yeast *Saccharomyces cerevisiae,* strain VL6-48N (MATαhis3-Δ200 trp1-Δ1 KlURA3-Δ1 lys2 ade2-101 met14) containing the 1.08 Mb genome of *Mycoplasma mycoides* subsp. *capri* (*Mmc*) strain GM12 with an integrated yeast centromeric plasmid (YCp) (13) was used for construction of the mutants. Yeast cells were grown and maintained in either synthetic minimal medium containing dextrose (SD, Takara Bio) (13), or in standard rich medium containing glucose (YPD, Takara Bio) or galactose (YPG, Takara Bio) (32). SD medium was supplemented with 5-fluoroorotic acid (5-FOA), for KlURA3 counter-selection (13, 33).

### Preparation of mutagenesis cassettes

Sixty eight genes that encode candidate virulence traits were seamlessly deleted from the genome of *Mmc* GM12::YCpMmyc1.1 in five consecutive cycles (D1, D3, D4 and D5) using Tandem Repeat coupled with Endonuclease Cleavage [TREC] as described (32, 34) or a variation of TREC involving the Cre-lox system for the D2 deletion, see below. Primer sequences to target and confirm the insertion of the mutagenesis cassette into each target site and to verify seamless deletion of the targeted genes are shown in Table S 1.

The *gts* gene cluster (D2) was targeted and deleted in the *Mmc* genome in the back-ground of the *glpFKO* deletion strain by employing a derivative of the *Mmc* synthetic cell JCVI-syn1.0 (35). Primers RC0905 and RC0906 (Table S1) were used to amplify the mutagenesis cassette from the synthetic cell derivative and was targeted to the *gts* region. Specific primers were used to confirm correct insertion at the target site by amplifying the junctions between the GM12::YCpMmyc1.1 genome and the inserted cassette. Galactose induction resulted in the Cre-mediated deletion of the *gts* region, leaving 13 bp of the 5’ end of the *gtsA* region, the 34 bp *loxP* site, and 27 bp of the 3’ end of the *lppB* gene. Specific primers were used to verify the knock-out.

### Transformation and PCR analysis

Transformation of the CORE3 cassette was performed by the lithium acetate method as described previously (36). Transformed yeast were plated on appropriate selection media [SD medium minus His (Teknova, CA) or SD medium (minus His and minus Ura)] and incubated at 30°C for 48 hours. Yeast colonies were patched on appropriate selective media and total DNA was isolated for PCR screening (37). The correct insertion of the mutagenesis cassette was verified by PCR amplification using upstream and downstream specific primers (Integrated DNA Technologies, Coralville, IA, USA) (Table S1).

### Transplantation

The modified GM12::YCpMmyc1.1 genomes (D1 to D5) were transplanted into *Mycoplasma capricolum* subsp. *capricolum* (*Mcc*) recipient cells with polyethylene glycol and selected for tetracycline resistance as described previously (13, 37). The resulting mutant strains were subjected to multiplex PCRs and pulsed-field gel electrophoresis as described elsewhere (38) to confirm integrity of the genome.

### Confirmation of the mutants using next generation sequencing and mapping assembly

Total DNA of the strains GM12, GM12::YCpMmyc1.1 and GM12::YCpMmyc1.1-Δ*68* was isolated as described before (39). DNA was sheared using sonication and subjected to Illumina sequencing using a MiSeq machine by University of California Santa Cruz, CA (USA). Reads were mapped to the designed genome sequences based on the parental strains GM12 and GM12::YCpMmyc1.1 and GM12::YCpMmyc1.1-Δ*68*. The raw reads (300bp PE) were QC with FastQC (https://www.bioinformatics.babraham.ac.uk/projects/fastqc/). The corrected reads were mapped onto the reference genome WT-YCP.fa with bwa mem (40) and converted to sorted bam with samtools (41). The bam files were analyzed for deletions using Delly2 (42) and Sprites (43), and the predictions validated visually using IGV (44). The list of strains and their deleted regions is summarized in Table 2.

**Table 2:**
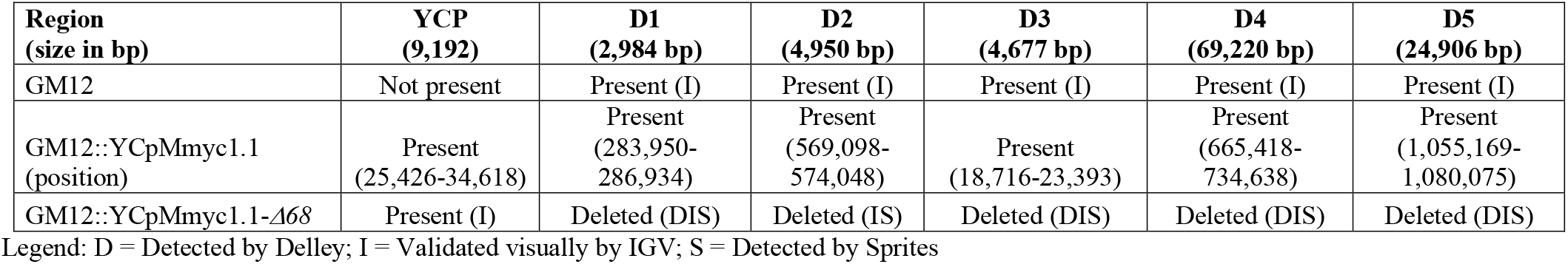
Table showing the results of the Illumina sequencing-based mapping assemble of the strains used in this study.

| **Region**<br>**(size in bp)** | **YCP**<br>**(9,192)** | **D1**<br>**(2,984 bp)** | **D2**<br>**(4,950 bp)** | **D3**<br>**(4,677 bp)** | **D4**<br>**(69,220 bp)** | **D5**<br>**(24,906 bp)** |
| --- | --- | --- | --- | --- | --- | --- |
| GM12 | Not present | Present (I) | Present (I) | Present (I) | Present (I) | Present (I) |
| GM12::YCpMmyc1.1<br>(position) | Present<br>(25,426-34,618) | Present<br>(283,950-286,934) | Present<br>(569,098-574,048) | Present<br>(18,716-23,393) | Present<br>(665,418-734,638) | Present<br>(1,055,169-1,080,075) |
| GM12::YCpMmyc1.1-Δ68 | Present (I) | Deleted (DIS) | Deleted (IS) | Deleted (DIS) | Deleted (DIS) | Deleted (DIS) |

Legend: D = Detected by Delley; I = Validated visually by IGV; S = Detected by Sprites

### Scanning electron microscopy of *Mycoplasma*

Unless stated otherwise, chemicals were obtained from Merck (Schaffhausen, Switzerland). Mycoplasmas were washed with distilled water (dH_2_O) and fixed with 4% para-formaldehyde (Life Technologies, Thermo Fisher, Zug, Switzerland; Cat. No. 28906) in dH_2_O for five days at 4°C. Thereafter, samples of 40 μl of cell suspension were centrifuged onto gold-sputtered poly-L-lysine coated coverslips (high molecular poly-L-Lysine hydrobromide) at 125 *rcf* for 4 min. Coverslips were washed 1x with PBS and 2x with 0.1% bovine serum albumin in PBS (BSA/PBS). Free aldehydes were blocked with 0.05M glycine in 0.1% BSA-c/PBS (Aurion, ANAWA Trading, Wangen, Switzerland) for 15 min. at room temperature. After 3x washes with 0.1% BSA/PBS, cells were fixed with 2.5% glutaraldehyde (Merck 104239) in 0.1M cacodylate buffer (dimethylarsinic acid sodium salt trihydrate), washed 3x with dH_2_O and postfixed with 1% OsO_4_ (Polysciences, Warrington, PA, USA) in 0.1M cacodylate buffer for 15 min. at room temperature. Another 5x washes with dH_2_O were followed by dehydration in an ascending ethanol series. Samples were then transferred to hexamethyldisilazane (Merck 814051) for 10 min, air-dried, mounted onto aluminum stubs with carbon conductive adhesive tabs (Ted Pella Inc., Redding, CA, USA) and coated with approximately 25 nm of gold in an SCD004 (Leica Microsystems, Heerbrugg, Switzerland). Secondary electron micrographs and corresponding backscattered images were obtained with a fully digital field emission scanning electron microscope DSM 982 Gemini (Zeiss, Oberkochen, Germany) at an accelerating voltage of 5 kV, a working distance of 6–8 mm and primary magnifications ranging from 30,000 to 50,000×.

### Growth assay

Overnight cultures of *Mycoplasma* strains were grown at 37°C in SP4 medium containing streptomycin (GM12) or tetracycline (GM12::YCpMmyc1.1, GM12::YCpMmyc1.1-*∆68*) for about 16 hours. Doubling times of the Mycoplasma strains were then determined as described elsewhere (24), except that time interval samples were collected and processed at 0, 1, 2, 3, 4, 5, 6, 7, 9, 12, 15, and 24 h.

### *In vitro* hydrogen peroxide assay

Overnight cultures of *Mycoplasma* strains were grown as described above. When the pH of overnight cultures reached 6.0 - 6.5, they were inoculated into fresh SP4 medium at 1:200 dilution and incubated at 37 °C for different time intervals of 0, 5, 7, and 24 h. At each time interval, an aliquot of culture was taken for DNA extraction (24) and another aliquot was taken to determine hydrogen peroxide levels.

To determine hydrogen peroxide levels, the aliquots were spun at 14,000 rpm for 10 minutes at 4°C. The pellets were washed with 1 ml of cold PBS, pH 7.5 to remove traces of media, then resuspended in 400 μl of cold PBS and stored at 4°C. Hydrogen peroxide levels were determined using the Amplex Red Hydrogen Peroxide Assay Kit (Life Technologies, NY) according to the manufacturer’s instructions. Briefly, 50 μl of diluted samples (1:5 in PBS) was aliquoted onto 96-well plates and warmed to 37°C for 1 h prior to starting the assay. 100 μM final concentration of glycerol (Sigma Aldrich, MO) or GPC (Sigma Aldrich, MO) was then added to the diluted sample and incubated at 37°C for 1 h. 50 μl of the Amplex Red reagent was added to the samples, incubated at room temperature in the dark for 30 minutes and fluorescence was measured using a spectrophotometer (SpectraMax M5, Molecular Devices, CA). Three technical replicates were performed for each sample and normalized to their respective DNA concentrations.

### Detection of immunoglobulin degradation by *Mycoplasma in vitro* and *in vivo*

*In vitro* functionality of the MIP-MIP system was tested using the strains GM12, GM12::YCpMmyc1.1 and GM12::YCpMmyc1.1-Δ*68*. Each strain was grown overnight at 37°C in 3 mL modified SP5 medium (containing 5% FBS). 250 μL of each culture was harvested and centrifuged for 10 minutes at 4,000 *g*. The cells were then resuspended in 15 μL modified SP5 medium (containing 5% FBS) and incubated with 5 μg of purified caprine IgG (Sigma) at 37°C for 3 hours. Bacterial CFUs were estimated for each strain by serial dilutions and were 3.2*10^9^ CFUs *Mmc* GM12, 8.5*10^8^ CFUs *Mmc* GM12::YCpMmyc1.1 and 3.5*10^8^ CFUs *Mmc* GM12::YCpMmyc1.1-Δ*68*. A sample consisting of 5 μg caprine IgG in dH_2_O was included as a control. The incubated samples were mixed with 2x Laemmli Sample Buffer (BioRad,) at a 1:1 ratio, boiled for 10 minutes at 98°C and separated onto a 12% SDS-PAGE gel. They were subsequently transferred onto a 0.2 μm nitrocellulose membrane (Bio-Rad) using a Bio-Rad Trans-Blot® Turbo^™^ Transfer System (25 volts, 1.0 A, 30 minutes). Next, a Western Blot was performed using PBS supplemented with 0.1% Tween-20 and 2% BSA as a blocking buffer, and mouse anti-goat IgG (H+L) (Jackson Immuno Research, 205-005-108) and goat anti-mouse (Fc) labelled with horseradish peroxidase (Sigma, A0168) as primary and secondary antibodies. The antibodies were diluted in blocking buffer at 1:2000 and 1:70,000, respectively, and incubated with the membrane for 1 hour each. In between the antibody incubations, the membrane was washed once with PBS – 0.1% Tween-20 + 3.2% NaCl and twice with PBS – 0.1% Tween-20, for 10 minutes each time. The results were visualized using the Fujifilm LAS-3000 Luminescent Image Analyzer.

Serum samples derived from the *in vivo* challenge were diluted in water to achieve a total load of 10-20 μg of protein per sample. Western Blots were performed and analyzed using the same protocol as described above.

### Animal experiment setup

All protocols of this study were designed and performed in strict accordance with the Kenyan and US American legislation for animal experimentation and were approved by the institutional animal care and use committee at both institutions (JCVI and ILRI, IACUC reference number 2014.08).

Sixteen male outbred goats (*Capra aegagrus hircus*), 1-2 years of age and randomly selected in Naivasha, were transferred to the ILRI campus in Nairobi and kept under quarantine for 6 months. After arrival at the campus, all animals were dewormed twice using levamisole and treated prophylactically against babesiosis and anaplasmosis using imidocarb. Upon entry to ILRI, the goats were vaccinated against anthrax & blackleg (Blanthax®, Cooper), Foot and Mouth Disease (FOTIVAX®) and Peste des Petits Ruminants (Live attenuated strain Nig. 75/1). All animals were tested negative for presence of antibodies against CCPP, using a competitive ELISA (IDEXX). Two weeks before experimental infection, all animals were transferred to the animal biosafety level two (ABSL2) unit. *Mycoplasma* cells were cultivated in PPLO medium supplemented with horse serum (45) to early logarithmic phase, aliquoted and stored at −80°C. Afterwards, we determined the CFU using two aliquots. Just before infection we thawed the vials and adjusted the concentration of *Mycoplasma* to 10^9^CFU per mL^-1^ using broth. All 16 goats were infected transtracheally by needle puncture 5-10 cm distal to the larynx. Each animal received 1 mL of *Mmc* GM12 or GM12::YCpMmyc1.1-Δ*68* liquid culture (10^9^ colony forming units per animal), followed by 5 mL of phosphate buffered saline (PBS). The animals were allowed to move freely within the ABSL2 unit and had *ad libitum* access to water. They were fed *ad libitum* with hay and received pellets each morning. Three veterinarians monitored the health status of the animals throughout the experiment. Rectal temperature, oxygen blood saturation, heart rate & breathing frequency were measured daily in the morning hours using the GLA M750 thermometer (GLA agricultural electronics, USA), VE-H100B oximeter (Edan, USA), and a stethoscope classic II (Littmann, USA) with a water-resistant wrist watch Seamaster (Omega, Switzerland), respectively. Blood samples for subsequent analysis were taken twice a week by jugular vein puncture. Nasal swabs (Flocked swabs, Copan, Italy) were taken twice a week. Swabs were transferred into cryotubes filled with media and stored at −80°C until further processing. Goats were euthanized when they developed severe disease associated with unwarranted moderate to severe pain. Therefore, they received an intravenous injection of Lethabarb Euthanasia Injection (Virbac, USA) of 200 mg.kg−^1^ body weight. Severe disease and pain were determined by a fever of ≥41°C for >3 consecutive days, an oxygen saturation of ≤92% and a lateral recumbency of ≥1 day without the ability to feed or intake water. Goats that were not put down because of ethical reasons were euthanized on 28 dpi.

### Pathomorphological and histology analysis

A complete necropsy was performed on all animals. Tissue samples of the neck region around the inoculation site and all internal organs were fixed in 10 % buffered formalin for 72 hours and subsequently routinely processed for paraffin embedding. Tissue sections were cut at 3 μm and stained routinely with hematoxylin and eosin (H&E) and evaluated by a board-certified pathologist.

### Microbiology

Venous blood samples, lung samples, carpal joint fluid, and pleural fluid specimens taken at necropsy were used for isolation of *Mmc* as described elsewhere (46) using *Mycoplasma* liquid medium (Mycoplasma Experience, Ltd., United Kingdom). Lung samples and pleural fluid were used for screening of *Pasteurella* and *Mannheimia* spp. using standard methods (47).

### Statistical analysis

Exact and normal approximation binomial tests were used to compare the two groups using GenStat 12th Edition (48). P values for differences in parameters were estimated using a 2sided 2-sample t-test comparing average levels between both groups at 5% level of significance.

## ACKNOWLEDGMENTS

The National Science Foundation [grant number IOS-1110151 (to S.V., C.L., and J.J.)] provided the funds for this work. Additional support was received from the University of Bern and the CGIAR research program on Livestock and Fish. Elise Schieck was supported by the German Federal Ministry for Economic Cooperation and Development (Project No. 09.7860.1-001.00 Contract no. 81136800, http://www.bmz.de). Anne Liljander was supported by the Centrum for International Migration and Development in Germany. We thank Ray-Yuan Chuang, Vladimir Noskov, Steven Weber, Pamela Nicholson, Helga Mogel, Elisabeth Cook and the ILRI animal caretakers for their excellent advice and help.

## AUTHOR CONTRIBUTIONS

J.J. and S.V. designed research; L.M., N.G-A., S.C., J.J., A.L., P.S., M.S., E.S., V.C., Y.A., HP. and F.L. performed research; J.J., F.L., S.C., Y.A., L.F., P.S-P., C.L., A.B. and SV., analyzed data; J.J. and S.V. wrote the paper

---

**Table S1:**
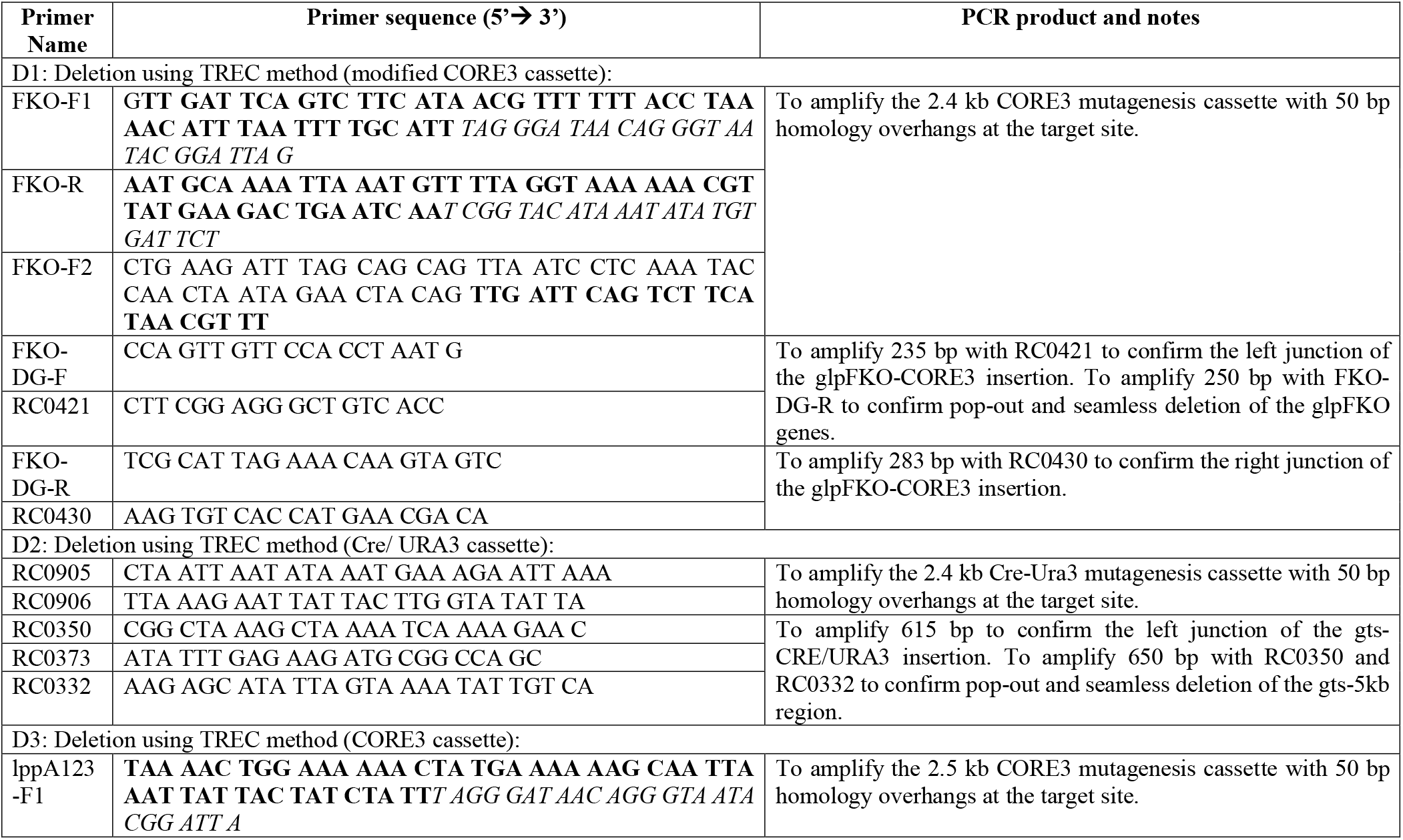
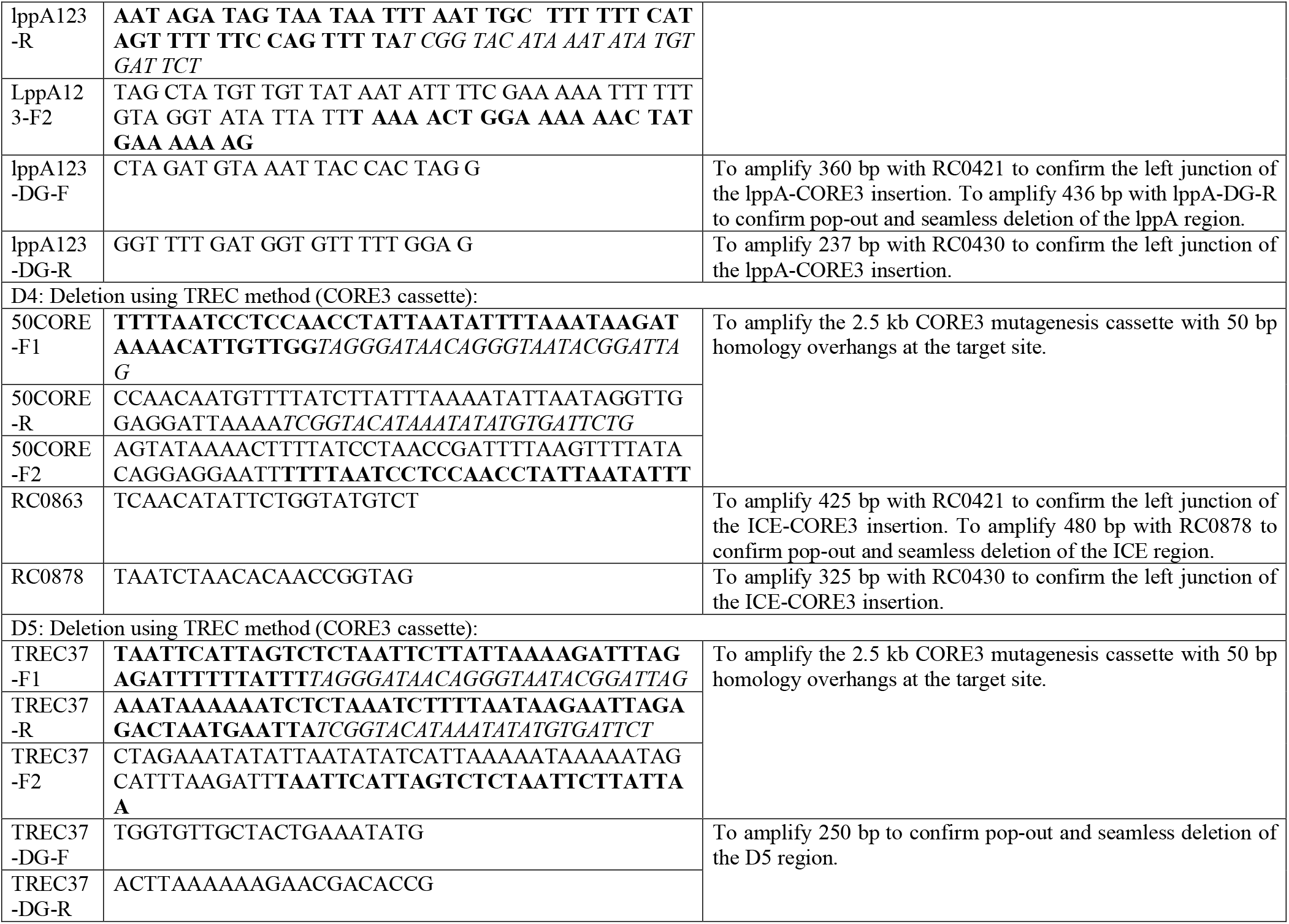
Oligonucleotide primers used to construct the mutagenesis cassettes and to confirm the deletions.

**Figure S1:**
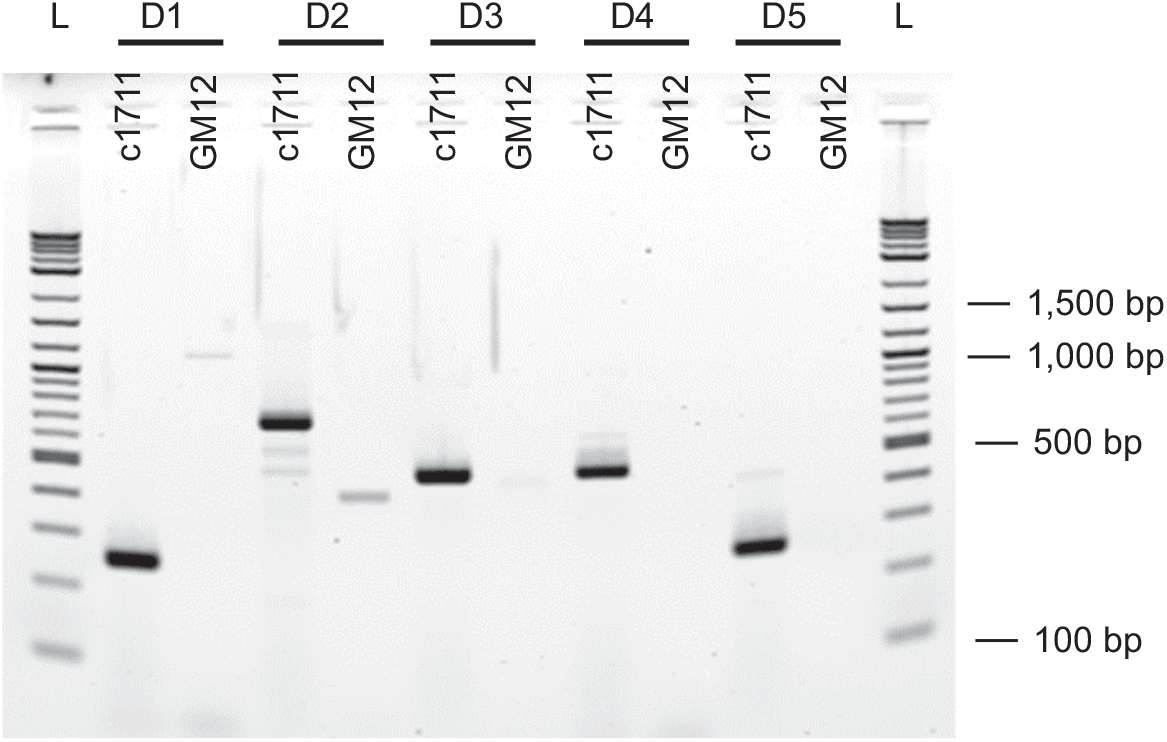
Specific PCRs confirming the deletions in GM12::YCpMmyc1.1-*∆68*. Genomic DNA from GM12::YCpMmyc1.1-*∆68* (clone 1711) and its parental strain GM12 was used as template. Specific primers were used to amplify across the deleted regions. DNA from GM12::YCpMmyc1.1-*∆68* (clone 1711) yielded expected amplicons that are a product covering the flanking regions of the deleted subgenomic fragment. Diagnostic primers FKO-DG-F and FKO-DG-R result in a 250 bp amplicon across the D1 region in clone 1711, primers RC0350 and RC0332 result in a 650 bp amplicon across the D2 region, primers LppA123-DG-F and LppA123-DG-R result in a 436 bp amplicon across the D3 region, primers RC0863 and RC0878 result in a 480 bp amplicon across the D4 region, and primers TREC37-DG-F and TREC37-DG-R result in a 250 bp region across the D5 region. L denotes the DNA ladder.

## REFERENCES

1. Linchevski I, Klement E, Nir-Paz R. 2009. *Mycoplasma pneumoniae* vaccine protective efficacy and adverse reactions--Systematic review and meta-analysis. Vaccine 27:2437–46.

2. Browning GF, Whithear KG, Geary SJ. 2005. Vaccines to control Mycoplasmosis, p 569–597. In Blanchard A, Browning G (ed), Mycoplasmas : molecular biology, pathogenicity and strategies for control. Horizon Bioscience, Wymondham, Norfolk, U.S.A.

3. Jores J, Mariner JC, Naessens J. 2013. Development of an improved vaccine for contagious bovine pleuropneumonia: an African perspective on challenges and proposed actions. Vet Res 44:122.

4. Nicholas R, Churchward C. 2012. Contagious caprine pleuropneumonia: new aspects of an old disease. Transbound Emerg Dis 59:189–96.

5. Maes D, Segales J, Meyns T, Sibila M, Pieters M, Haesebrouck F. 2008. Control of *Mycoplasma hyopneumoniae* infections in pigs. Veterinary microbiology 126:297–309.

6. Chopra-Dewasthaly R, Spergser J, Zimmermann M, Citti C, Jechlinger W, Rosengarten R. 2017. Vpma phase variation is important for survival and persistence of Mycoplasma agalactiae in the immunocompetent host. PLoS Pathog 13:e1006656.

7. Citti C, Nouvel LX, Baranowski E. 2010. Phase and antigenic variation in mycoplasmas. Future Microbiol 5:1073–85.

8. Chambaud I, Wroblewski H, Blanchard A. 1999. Interactions between mycoplasma lipoproteins and the host immune system. Trends Microbiol 7:493–499.

9. Arfi Y, Minder L, Di Primo C, Le Roy A, Ebel C, Coquet L, Claverol S, Vashee S, Jores J, Blanchard A, Sirand-Pugnet P. 2016. MIB-MIP is a mycoplasma system that captures and cleaves immunoglobulin G. Proc Natl Acad Sci U S A 113:5406–5411.

10. Blotz C, Stulke J. 2017. Glycerol metabolism and its implication in virulence in *Mycoplasma*. FEMS Microbiol Rev 41:640–652.

11. Pilo P, Frey J, Vilei EM. 2007. Molecular mechanisms of pathogenicity of *Mycoplasma mycoides* subsp. *mycoides* SC. Vet J 174:513–21.

12. Szczepanek SM, Boccaccio M, Pflaum K, Liao X, Geary SJ. 2014. Hydrogen peroxide production from glycerol metabolism is dispensable for virulence of *Mycoplasma gallisepticum* in the tracheas of chickens. Infect Immun 82:4915–20.

13. Lartigue C, Vashee S, Algire MA, Chuang RY, Benders GA, Ma L, Noskov VN, Denisova EA, Gibson DG, Assad-Garcia N, Alperovich N, Thomas DW, Merryman C, Hutchison CA, 3rd, Smith HO, Venter JC, Glass JI. 2009. Creating bacterial strains from genomes that have been cloned and engineered in yeast. Science 325:1693–6.

14. Browning GF, Marenda MS, Noormohammadi AH, Markham PF. 2011. The central role of lipoproteins in the pathogenesis of mycoplasmoses. Veterinary microbiology 153:44–50.

15. Beven L, Charenton C, Dautant A, Bouyssou G, Labroussaa F, Skollermo A, Persson A, Blanchard A, Sirand-Pugnet P. 2012. Specific evolution of F1-like ATPases in mycoplasmas. PLoS One 7:e38793.

16. Tardy F, Mick V, Dordet-Frisoni E, Marenda MS, Sirand-Pugnet P, Blanchard A, Citti C. 2015. Integrative conjugative elements are widespread in field isolates of *Mycoplasma* species pathogenic for ruminants. Appl Environ Microbiol 81:1634–43.

17. Krasteva I, Liljander A, Fischer A, Smith DG, Inglis NF, Scacchia M, Pini A, Jores J, Sacchini F. 2014. Characterization of the *in vitro* core surface proteome of *Mycoplasma mycoides* subsp. *mycoides*, the causative agent of contagious bovine pleuropneumonia. Vet Microbiol 168:116–23.

18. Weldearegay YB, Pich A, Schieck E, Liljander A, Gicheru N, Wesonga H, Thiaucourt F, Kiirika LM, Valentin-Weigand P, Jores J, Meens J. 2016. Proteomic characterization of pleural effusion, a specific host niche of *Mycoplasma mycoides* subsp. *mycoides* from cattle with contagious bovine pleuropneumonia (CBPP). J Proteomics 131:93–103.

19. Jores J, Schieck E, Liljander A, Sacchini F, Posthaus H, Lartigue C, Blanchard A, Labroussaa F, Vashee S. 2018. Capsular polysaccharide in *Mycoplasma mycoides* is a virulence factor. J Infect Dis doi:10.1093/infdis/jiy713.

20. Falkow S. 1988. Molecular Koch’s postulates applied to microbial pathogenicity. Rev Infect Dis 10 Suppl 2:S274–6.

21. Monnerat MP, Thiaucourt F, Poveda JB, Nicolet J, Frey J. 1999. Genetic and serological analysis of lipoprotein LppA in *Mycoplasma mycoides* subsp. *mycoides* LC and *Mycoplasma mycoides* subsp. *capri*. Clin Diagn Lab Immunol 6:224–30.

22. Dedieu L, Totte P, Rodrigues V, Vilei EM, Frey J. 2010. Comparative analysis of four lipoproteins from *Mycoplasma mycoides* subsp. *mycoides* Small Colony identifies LppA as a major T-cell antigen. Comp Immunol Microbiol Infect Dis 33:279–90.

23. Mulongo M, Frey J, Smith K, Schnier C, Wesonga H, Naessens J, McKeever D. 2015. Vaccination of cattle with the N terminus of LppQ of *Mycoplasma mycoides* subsp. *mycoides* results in type III immune complex disease upon experimental infection. Infection and immunity 83:1992–2000.

24. Hutchison CA, 3rd, Chuang RY, Noskov VN, Assad-Garcia N, Deerinck TJ, Ellisman MH, Gill J, Kannan K, Karas BJ, Ma L, Pelletier JF, Qi ZQ, Richter RA, Strychalski EA, Sun L, Suzuki Y, Tsvetanova B, Wise KS, Smith HO, Glass JI, Merryman C, Gibson DG, Venter JC. 2016. Design and synthesis of a minimal bacterial genome. Science 351:aad6253.

25. Schieck E, Lartigue C, Frey J, Vozza N, Hegermann J, Miller RA, Valguarnera E, Muriuki C, Meens J, Nene V, Naessens J, Weber J, Lowary TL, Vashee S, Feldman MF, Jores J. 2016. Galactofuranose in *Mycoplasma mycoides* is important for membrane integrity and conceals adhesins but does not contribute to serum resistance. Mol Microbiol 99:55–70.

26. Richter SH, Garner JP, Auer C, Kunert J, Wurbel H. 2010. Systematic variation improves reproducibility of animal experiments. Nat Methods 7:167–8.

27. Mogensen TH. 2009. Pathogen recognition and inflammatory signaling in innate immune defenses. Clin Microbiol Rev 22:240–73.

28. Hirayasu K, Saito F, Suenaga T, Shida K, Arase N, Oikawa K, Yamaoka T, Murota H, Chibana H, Nakagawa I, Kubori T, Nagai H, Nakamaru Y, Katayama I, Colonna M, Arase H. 2016. Microbially cleaved immunoglobulins are sensed by the innate immune receptor LILRA2. Nat Microbiol 1:16054.

29. Abu Kwaik Y, Bumann D. 2013. Microbial quest for food *in vivo*: ‘nutritional virulence’ as an emerging paradigm. Cell Microbiol 15:882–90.

30. Fischer A, Shapiro B, Muriuki C, Heller M, Schnee C, Bongcam-Rudloff E, Vilei EM, Frey J, Jores J. 2012. The Origin of the ‘*Mycoplasma mycoides* Cluster’ Coincides with Domestication of Ruminants. PLoS One 7:e36150.

31. DaMassa AJ, Brooks DL, Adler HE. 1983. Caprine mycoplasmosis: widespread infection in goats with *Mycoplasma mycoides* subsp *mycoides* (large-colony type). American journal of veterinary research 44:322–5.

32. Noskov VN, Segall-Shapiro TH, Chuang RY. 2010. Tandem repeat coupled with endonuclease cleavage (TREC): a seamless modification tool for genome engineering in yeast. Nucleic acids research 38:2570–6.

33. Boeke JD, LaCroute F, Fink GR. 1984. A positive selection for mutants lacking orotidine-5’-phosphate decarboxylase activity in yeast: 5-fluoro-orotic acid resistance. Mol Gen Genet 197:345–6.

34. Chandran S, Noskov VN, Segall-Shapiro TH, Ma L, Whiteis C, Lartigue C, Jores J, Vashee S, Chuang R. 2014. TREC-IN: gene knock-in genetic tool for genomes cloned in yeast. BMC Genomics 15:1180.

35. Gibson DG, Glass JI, Lartigue C, Noskov VN, Chuang RY, Algire MA, Benders GA, Montague MG, Ma L, Moodie MM, Merryman C, Vashee S, Krishnakumar R, Assad-Garcia N, Andrews-Pfannkoch C, Denisova EA, Young L, Qi ZQ, Segall-Shapiro TH, Calvey CH, Parmar PP, Hutchison CA, 3rd, Smith HO, Venter JC. 2010. Creation of a Bacterial Cell Controlled by a Chemically Synthesized Genome. Science 329:52–56.

36. Gietz D, St Jean A, Woods RA, Schiestl RH. 1992. Improved method for high efficiency transformation of intact yeast cells. Nucleic Acids Res 20:1425.

37. Noskov V, Kouprina N, Leem SH, Koriabine M, Barrett JC, Larionov V. 2002. A genetic system for direct selection of gene-positive clones during recombinational cloning in yeast. Nucleic Acids Res 30:E8.

38. Labroussaa F, Lebaudy A, Baby V, Gourgues G, Matteau D, Vashee S, Sirand-Pugnet P, Rodrigue S, Lartigue C. 2016. Impact of donor-recipient phylogenetic distance on bacterial genome transplantation. Nucleic Acids Res 44:8501–11.

39. Fischer A, Santana-Cruz I, Hegerman J, Gourle H, Schieck E, Lambert M, Nadendla S, Wesonga H, Miller RA, Vashee S, Weber J, Meens J, Frey J, Jores J. 2015. High quality draft genomes of the *Mycoplasma mycoides* subsp. *mycoides* challenge strains Afadé and B237. Stand Genomic Sci 10:89.

40. Li H, Durbin R. 2010. Fast and accurate long-read alignment with Burrows-Wheeler transform. Bioinformatics 26:589–95.

41. Li H. 2011. Improving SNP discovery by base alignment quality. Bioinformatics 27:1157–8.

42. Rausch T, Zichner T, Schlattl A, Stutz AM, Benes V, Korbel JO. 2012. DELLY: structural variant discovery by integrated paired-end and split-read analysis. Bioinformatics 28:i333–i339.

43. Zhang Z, Wang J, Luo J, Ding X, Zhong J, Wang J, Wu FX, Pan Y. 2016. Sprites: detection of deletions from sequencing data by re-aligning split reads. Bioinformatics 32:1788–96.

44. Thorvaldsdottir H, Robinson JT, Mesirov JP. 2013. Integrative Genomics Viewer (IGV): high-performance genomics data visualization and exploration. Brief Bioinform 14:178–92.

45. Sacchini F, Naessens J, Awino E, Heller M, Hlinak A, Haider W, Sterner-Kock A, Jores J. 2011. A minor role of CD4+ T lymphocytes in the control of a primary infection of cattle with *Mycoplasma mycoides* subsp. *mycoides*. Vet Res 42:77.

46. Liljander A, Yu M, O’Brien E, Heller M, Nepper JF, Weibel DB, Gluecks I, Younan M, Frey J, Falquet L, Jores J. 2015. A field-applicable recombinase polymerase amplification assay for rapid detection of *Mycoplasma capricolum* subsp. *capripneumoniae*. J Clin Microbiol 53:2810–5.

47. Carter GR, Cole JR (ed). 1990. Diagnostic procedures in veterinary bacteriology and mycology. Academic Press, San Diego, Calif.

48. Payne RW, Murray DA, Harding SA, Baird DB, Soutar DM. 2012. GenStat for Windows (15th Edition) Introduction., VSN International, Hemel Hempstead, UK.

